# Small pangenome of *Candida parapsilosis* reflects overall low intraspecific diversity

**DOI:** 10.1101/2025.04.30.651475

**Authors:** Adam P. Ryan, Sean Bergin, Jillian Scully, Evelyn Zuniga-Soto, Conor Hession, Amelia E. Barber, Emilia Gomez-Molero, Oliver Bader, Florent Morio, Bing Zhai, Kenneth H. Wolfe, Geraldine Butler

## Abstract

*Candida parapsilosis* is an opportunistic yeast pathogen that can cause life-threatening infections in immunocompromised humans. Whole genome sequencing (WGS) studies of the species have demonstrated remarkably low diversity, with strains typically differing by about 1.5 single nucleotide polymorphisms (SNPs) per 10 kb. However, SNP calling alone does not capture the full extent of genetic variation. Here, we define the pangenome of 372 *C. parapsilosis* isolates to determine variation in gene content. The pangenome consists of 5,859 genes, of which 48 are not found in the genome of the reference strain. This includes 5,791 core genes (present in ≥ 99.5% of isolates). Four genes, including the allantoin permease gene *DAL4*, were present in all isolates but were truncated in some strains. The truncated *DAL4* was classified as a pseudogene in the reference strain CDC317. CRISPR-Cas9 gene editing showed that removing the early stop codon (producing the full-length Dal4 protein) is associated with improved use of allantoin as a sole nitrogen source. We find that the accessory genome of *C. parapsilosis* consists of 68 homologous clusters. This includes 38 previously annotated genes, 27 novel paralogs of previously annotated genes and 3 uncharacterised ORFs. Approximately one-third of the accessory genome (24/68 genes) is associated with gene fusions between tandem genes in the major facilitator superfamily (MFS). Additionally, we identified two highly divergent *C. parapsilosis* strains and find that, despite their increased phylogenetic distance (∼30 SNPs per 10 kb), both strains have similar gene content to the other 372.

**Importance:** *Candida parapsilosis* is a human fungal pathogen, listed in the high priority group by the World Health Organisation. It is an increasing cause of hospital-acquired and drug-resistant infection. Here, we studied the genetic diversity of 372 *C. parapsilosis* isolates, the largest genomic surveillance of this species to date. We show that there is relatively little genetic variation. However, we identified two more distantly-related isolates from Germany, suggesting that even more sampling may yield more diversity. We find that the pangenome (the cumulative gene content of all isolates) is surprisingly small, compared to other fungal species. Many of the non-core genes are involved in transport. We also find that variations in gene content are associated with nitrogen metabolism, which may contribute to the virulence characteristics of this species.

## Introduction

*C. parapsilosis* is diploid and belongs to the Serinales order (1–3). These yeasts translate the codon CUG as serine instead of leucine (2). *C. parapsilosis* infections can be superficial and invasive, and may arise from previous intestinal colonisation in immunocompromised individuals (4). *C. parapsilosis* infections are becoming increasingly prevalent globally, currently ranking in the top three of the most common *Candida* infections (5, 6). In particular, the number of outbreaks involving nosocomial transmission of fluconazole resistant isolates is increasing, posing serious threats to the health of immunocompromised patients (7). Prevalence differs by geography, with a high incidence in Southern Europe, South America, Asia and the Middle East, ranging from 11 to 41% by locality (5).

Previous analyses of genetic variation in *C. parapsilosis* have focused on small variants, SNPs and indels, and how they relate to health-related phenotypes such as antifungal resistance. Despite the overall low SNP diversity present in *C. parapsilosis* strains (8), these studies have successfully identified variants present in antifungal targets (9–11), multi-drug transporters (12), and regulators (10, 13) that contribute to resistance to the commonly used antifungal fluconazole. The high prevalence of some of these variants likely arises due to selective pressure from common use of fluconazole in clinical settings globally. Other work has focused on gene amplification and gene family expansion as sources of variation in *C. parapsilosis*. There is a broad variation in the copy number of the arsenite transporter *ARR3,* the phosphatidylcholine floppase *RTA3*, and *ALS* (adhesin) genes (14,15). Here we investigated a large set of 374 *C. parapsilosis* isolates, focusing on identifying variation in gene presence and absence through pangenome analysis.

Pangenome analysis assesses genetic variation by identifying orthologous gene clusters from multiple strains, and variation is catalogued as the presence and absence of clusters, or as changes in the numbers of clusters (16–19). Genes are typically split into three major groups: core, accessory and unique. The core genome contains genes found in all, or nearly all (e.g. >95%) samples. The accessory (or shell) genome contains genes found in only a portion of samples, typically applying an arbitrary minimum and maximum number (18, 20). The unique (or cloud) genome contains genes found in only one, or a defined small number, of samples (21). Unique genes may represent novel gene sequences, gene duplications that have since diverged, or genes acquired by horizontal gene transfer (HGT). They may also be artifacts resulting from failures in genome assembly or annotation.

Through *de novo* assembly, pangenomes are constructed by individually assembling and annotating each genome sequence (22, 23). Homologous gene clusters are identified and the pangenome is determined from the presence or absence of each sequence in each isolate. *De novo* assembly methods have been used to characterise the pangenomes of several yeast species. Analysis of *Saccharomyces cerevisiae* genomes identified 7,078 gene clusters across 1,392 isolates (24). On average 5,759 genes were found per isolate, and ∼75% of the total gene clusters were deemed to be core (24). In a similar study of 1,011 *S. cerevisiae* isolates, researchers found a pangenome size of 7,796 genes, with 4,940 core genes and 2,856 accessory genes (25). Because *de novo* assembly has had success in fungal species previously, we used this approach to construct and characterise the pangenome of *C. parapsilosis*, using a dataset of 374 sequenced isolates, including chromosome-level assemblies from at least one isolate from the five previously defined clades of this species (14).

## Results

### Diversity of genomes and phenotypes in a collection of 374 *C. parapsilosis* isolates

A SNP-based phylogeny of 374 *C. parapsilosis* isolates (including 64 previously described (14)) was constructed from short-read data aligned to the *C. parapsilosis* CDC317 reference genome. The majority of isolates were sourced from clinics in USA (29 isolates), France (45 isolates) and Germany (280 isolates) (Supplementary Table S1A). In brief, high quality biallelic SNPs were converted to FASTA format and heterozygous sites were randomly resolved on a site-by-site basis. The resulting FASTA alignment was used to generate a predicted phylogeny, with 1000 bootstrap replicates, using RAxML (26). A total of 96,501 informative sites were used to construct the tree (Figure 1).

**Figure 1.**
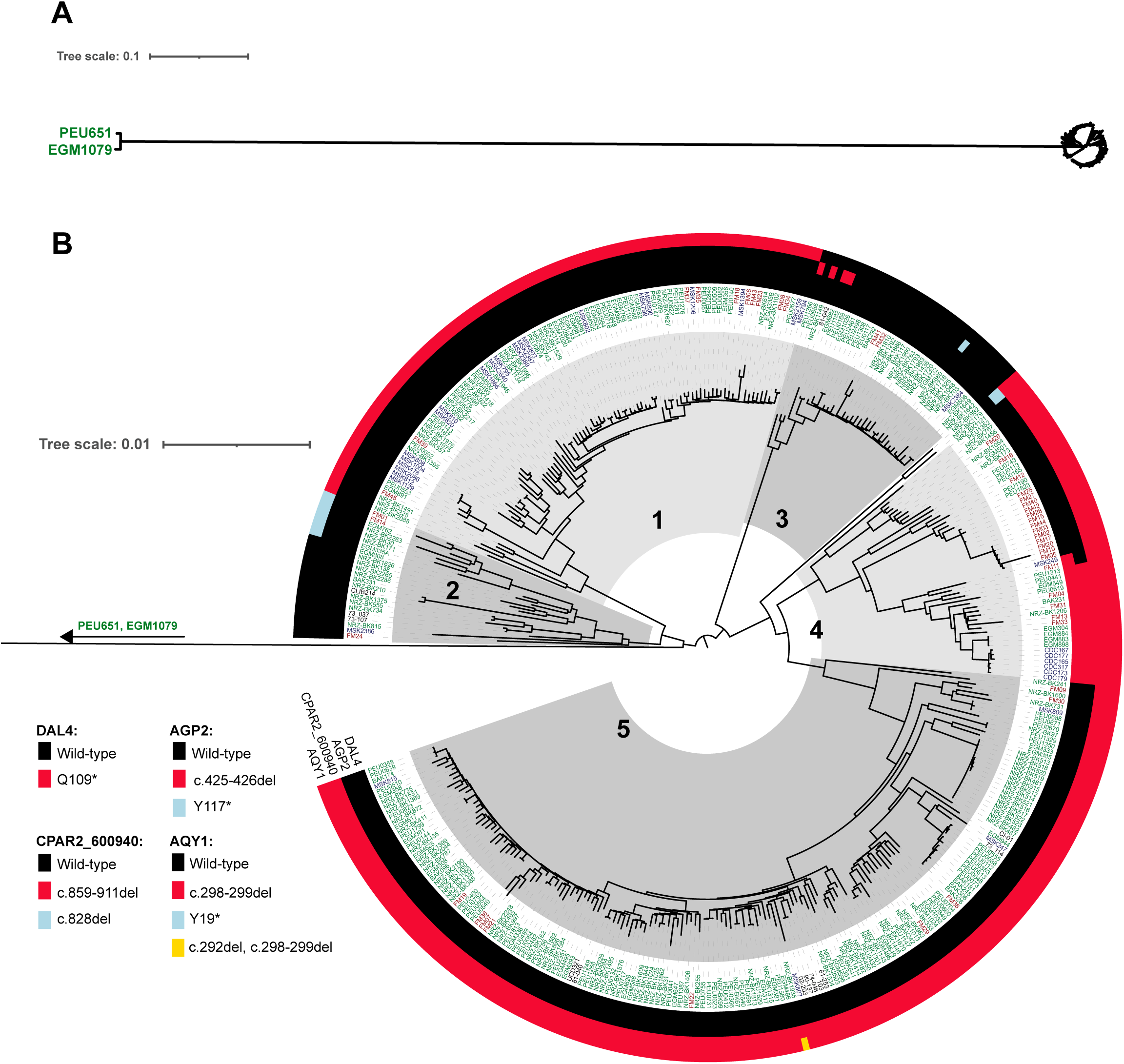
Estimated phylogeny of 374 *Candida parapsilosis* isolates, constructed using 96,501 SNP sites. (A) Full view of the tree including two distantly related isolates. (B) Zoomed-in view of the tree, cropping the long branch for clarity. The five major clades of *C. parapsilosis* are labeled. Strain names are coloured according to origin: France (red), Germany (green), USA (blue), and other (black). Outer rings indicate the presence of truncated isomorphs of four genes that cluster separately from the wild-type gene in the pangenome analysis. The corresponding colour for each allele in each gene is indicated on the left. From inner to outer, these are CPAR2_103200 (DAL4), CPAR2_401360 (AGP2), CPAR2_600940 and CPAR2_800150 (AQY1).

Isolates PEU651 and EGM1079 are unusually divergent from the other isolates and do not belong to the previously defined clades (Figure 1A). All but two of the other isolates fall into the five main clades described previously (14), with most (46%) belonging to Clade 5 (Figure 1B). Isolates NRZ-BK2179 and NRZ-BK982 lie slightly outside clades 4 and 5 (Figure 1B). Isolates show little geographic clustering, and every clade contains isolates from the USA, France, and Germany (Figure 1B). There are multiple examples where two strains from different countries are most closely related to each other, e.g. MSK249 (USA) and FM11 (France) in Clade 4.

The divergent isolates PEU651 and EGM1079 have 48,916 and 48,821 SNPs compared to the reference respectively, whereas the other 372 isolates have a mean of 2,656 SNPs (standard deviation of 647; Figure 2A). Interestingly, despite having approximately 20 times the total number of SNPs, PEU651 and EGM1079 have only twice as many heterozygous SNPs as the other isolates (Figure 2A). PEU651 and EGM1079 are still members of the species *C. parapsilosis*; they have ∼99.60% sequence identity to the reference genome, whereas all other strains have ∼99.98% identity. To prevent issues using divergent strains in identifying homologous gene clusters, we excluded PEU651 and EGM1079 from our systematic pangenome analysis, and instead inspected them separately manually.

**Figure 2.**
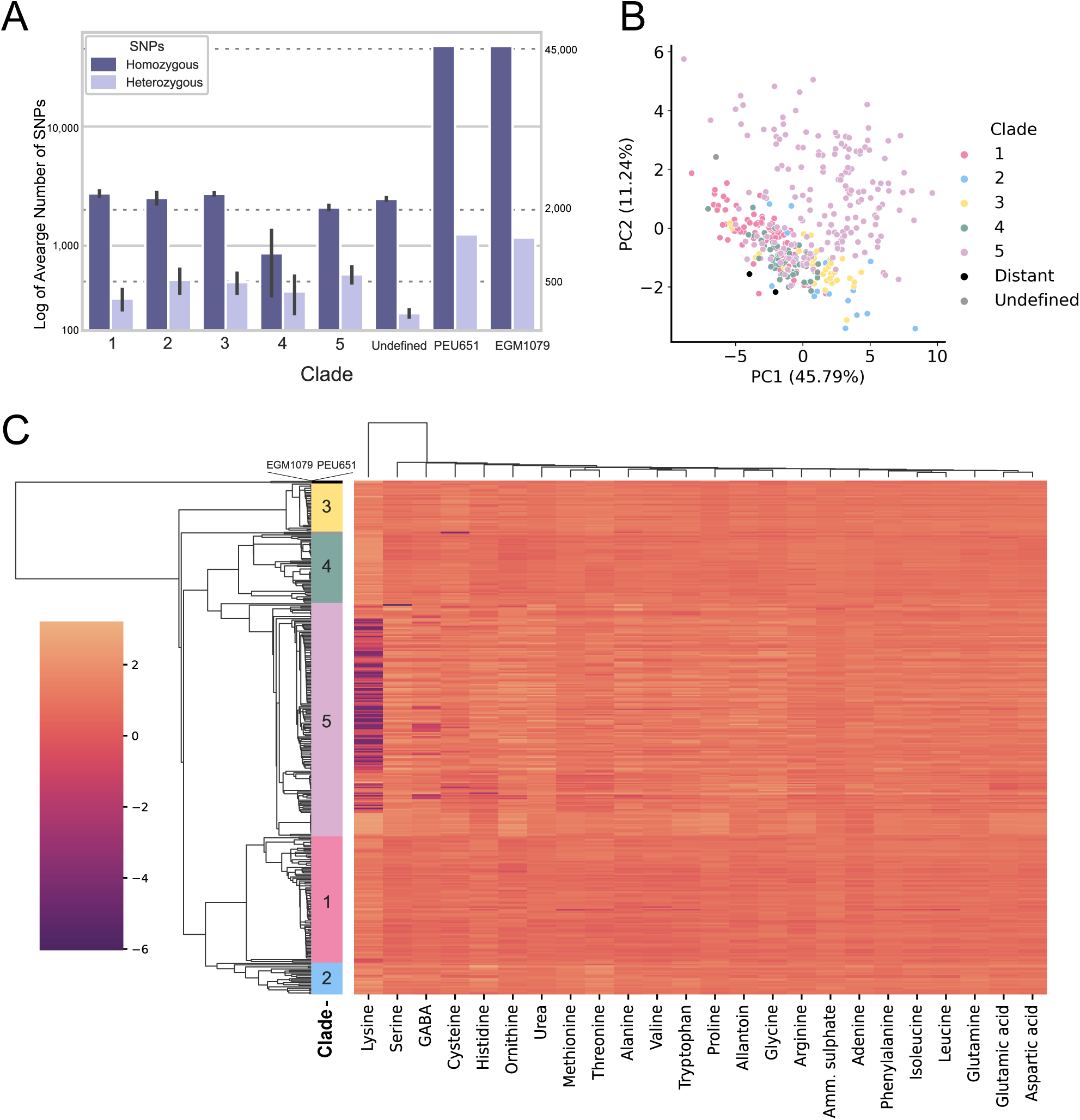
Isolates PEU651 and EGM1079 are genetically distant but phenotypically similar to the other *C. parapsilosis* isolates. (A) Mean numbers of homozygous and heterozygous SNPs identified relative to the CDC317 reference genome, shown on a log scale. PEU651 and EGM1079 are displayed separately, and other isolates are grouped by clade. Whiskers indicate the standard deviation of the mean. (B) Principal Components Analysis plot showing the first two principal components of the nitrogen-source phenotyping data. Each dot represents a single isolate coloured by clade. PC1 explains 45.78% of the variance of the dataset and PC2 explains 11.24%. (C) Clustered heatmap of growth of 374 strains (y-axis) on 24 nitrogen sources at 48h (x-axis). Darker colours represent reduced growth on nitrogen source when compared to the control plate. Strains are ordered using the same tree as in Figure 1, with the branch separating EGM1079 and PEU651 from the other strains shortened for visual clarity. Clustering of nitrogen sources was performed with the SciPy package, using the ‘average’ linking method with Euclidean distance.

Despite the increased phylogenetic distance of PEU651 and EGM1079 from the other 372 isolates, we observed little difference in their phenotypes. We measured growth rates of all 374 isolates on 23 different nitrogen sources (see Methods), and analysed the data by Principal Component Analysis (PCA). When the top two principal components are plotted, EGM1079 and PEU651 cluster near the majority of isolates (Figure 2B). Interestingly, strains from Clade 5 show the highest phenotypic diversity, spanning the breadth of both PC1 and PC2, whereas other clades show a tighter range overlapping each other (Figure 2B). This is likely due, at least in part, to the wide range of growth values on lysine rich conditions among Clade 5 isolates (Figure 2C).

### *De novo* assembly and annotation of *C. parapsilosis* isolate genomes

We used SPAdes (27) to generate genome assemblies of each isolate from the paired-end Illumina or DNB-seq short read sequencing data (Supplementary Table S1A). For nine genomes, chromosome-level assemblies were generated from long read (Oxford Nanopore) data using Canu (28), followed by error correction (polishing) with the short-read data (Supplementary Figure 1). Six of these genome assemblies were described previously (isolates MSK478, MSK802, MSK803, MSK812, CLIB214, and UCD321) (14, 29) and three more (PEU651, EGM1079, and NRZ-BK680) were sequenced here (Supplementary Table S1A). The genome sequence of the reference strain CDC317, which was sequenced by Sanger sequencing (1), was used as a chromosome-level assembly for Clade 4. Together, these data provide at least one chromosome-level assembly for each of the five main clades.

The chromosome-level assemblies had an average length of 13,086,549 ± 99,375 bp (Supplementary Table S1B). For short-read assemblies the average length was shorter, at 12,853,398 ± 62,418 bp with an average of 490 ± 97 contigs (Supplementary Table S1B). The genomes were annotated using BRAKER3 (30), trained using orthogroups derived from the Candida Gene Order Browser (CGOB) (31) (Supplementary Figures 1,2). The seven chromosome-level assemblies are predicted to encode 5,875 ± 46 genes, with a maximum of 5,957 in isolate UCD321 (Supplementary Table S1B). Similar numbers of genes were annotated in the short-read assemblies, with an average of 5,849 ± 46 genes per genome (Supplementary Table S1B). However, the maximum was much higher at 6,226 predicted genes for isolate NRZ-BK2265 (Supplementary Table S1B). In general, poorer assemblies (i.e. lower N50 values) correlated with higher gene counts (Supplementary Figure 3), suggesting that genome fragmentation was inflating the gene count. This is corroborated by counts of gene fragments annotated by BRAKER3. On average, BRAKER3 annotated 226 ± 52 fragmented gene sequences in scaffold-level assemblies but less than 1 per assembly at chromosome-level. The number of fragmented genes correlates with the number of contigs in scaffold-level assemblies (Supplementary Figure 3).

### The accessory genome of *C. parapsilosis* is small

BRAKER3 predicted 2,175,856 gene models across the 372 assemblies (workflows are shown in Supplementary Figures 1 and 2). Initial clustering of the gene models using GET_HOMOLOGS.pl (17) placed them into 7,241 homologous clusters, consisting of 6,067 non-unique clusters (from >2 isolates) and 1,174 unique clusters (from 1-2 isolates) (Supplementary Figure 1). Following manual merging and splitting of inconclusive clusters, the total number of homologous clusters was reduced from 6,067 to 5,785 (Supplementary Figure 2). These consisted of 5,703 single ortholog clusters, eight paired clusters, and 74 unresolved mixed clusters (Figure 3):

- The 5,703 single ortholog clusters represent 5,703 genes, of which 5,627 are present in the reference genome and 76 are novel.
- The eight paired clusters represent four variable-length genes that are truncated in some of the isolates. These genes are described in more detail below. They are all present in the reference genome.
- The 74 unresolved mixed clusters represent 152 genes (150 present in the reference and 2 novel paralogs) that could not be resolved to single orthologs because they comprised multiple copies of identical or near-identical genes, and gene fragments that could not be sorted into individual clusters. Coverage analysis showed that all 152 are present in at least one copy in every genome.

**Figure 3.**
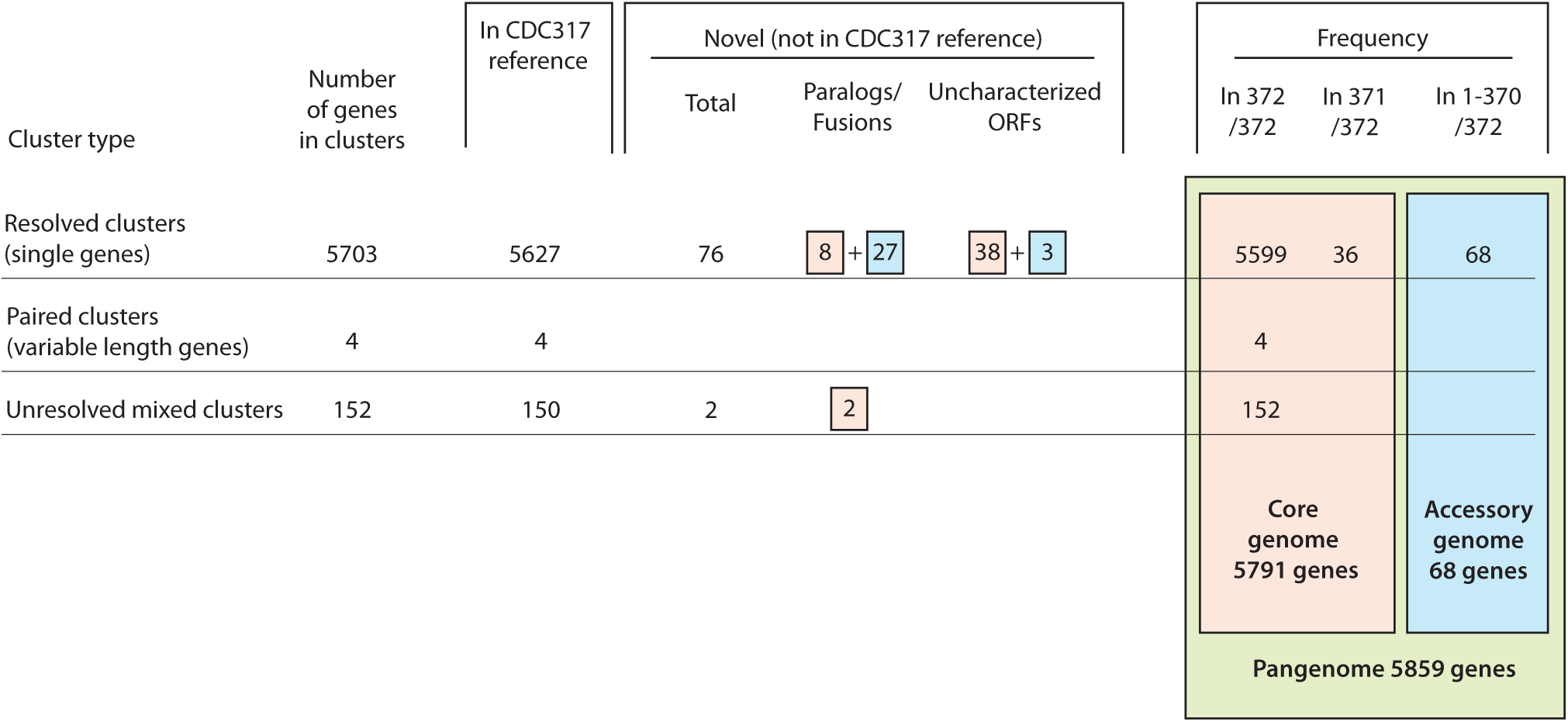
Pangenome analysis of *C. parapsilosis* pangenome. Genes highlighted in peach are assigned to the core genome, and those in blue are assigned to the accessory genome.

The pangenome of *C. parapsilosis* therefore consists of 5,859 genes, of which 5,703 are single resolved genes, 152 are unresolved genes, and 4 are paired (variable-length genes), as summarized in Figure 3. We used BLAST analysis to identify whether each of these genes is part of the core genome or is missing in individual isolates. All 152 unresolved genes and the four variable-length genes are found in all 372 isolates, and so are included in the core genome. Of the 5,703 resolved single genes, 5,599 were present in all 372 isolates and a further 36 were absent in only one isolate (Figure 3). These 5,635 genes were defined as part of the core genome. Among them, there are 46 novel genes not in the reference genome: 8 novel paralogs of genes previously annotated in CDC317, and 38 uncharacterised open reading frames (ORFs). In total, the core genome therefore contains 5,791 genes (Figure 3).

The accessory gene set comprises 68 single genes that are absent from at least two genomes, including 30 novel genes and 38 genes present in the reference (Supplementary Table S1C). The 30 novel accessory genes include 27 novel paralogs of reference genes (including fusions between genes) and 3 uncharacterised ORFs (Figure 3; Supplementary Table S1C). Ten of the identified novel genes overlap with novel transcriptional active regions (nTARs) annotated by Guida et al. (32), and two others are adjacent to nTARs, suggesting that they are transcribed.

Preliminary analysis indicated that presence or absence of accessory genes tends to be conserved in isolates that are related. To measure the strength of the phylogenetic signal of each accessory gene, Blomberg’s K-values were calculated (33, 34). Over half of the accessory genes (36 of 68) displayed a strong phylogenetic signal with high K-values (K ≥ 1, p < 0.05; Supplementary Table S1C), indicating a higher amount of similarity amongst related isolates than expected by chance (35). This observation suggests that gene loss or fusion often happened in ancestors of each clade.

### Four genes have truncated and non-truncated forms in different isolates

The four variable-length genes mentioned above correspond to four loci that each gave rise to two sequence clusters, corresponding to a long version and a short version of the protein, during automated sequence clustering (Table 1). For each of these genes, some isolates have an allele that generates a truncated or frameshifted protein, whereas all other isolates have an allele encoding a full-length protein, similar in length to its orthologs in other *Candida* species (31). For each of these loci, the CDC317 reference genome has a disrupted allele, resulting in shorter or frameshifted proteins (Table 1). Interestingly, two of these genes encode permeases, for pyrimidines (*DAL4*) and amino acids (*AGP2*), so the truncations could disrupt uptake of these molecules.

**Table 1.** Gene versions of paired homologous clusters harbouring premature stop codons. The allele present in the CDC317 reference sequence is indicated in bold.

| Gene | Variants (# isolates) | Standard name | InterPro description |
| --- | --- | --- | --- |
| *CPAR2_103200* | WT (351) <sup>1</sup><br>**C326T, Q109\* (21)** <sup>2</sup> | *DAL4* | Uracil/uridine/allantoin permease (IPR045225) [family] |
| *CPAR2_401360* | WT (350) <sup>1</sup><br>**c.425-426del (20)** <sup>3</sup><br>C351A, Y117* (2) <sup>2</sup> | *AGP2* | Amino Acid-Polyamine-Organocation Superfamily YAT (IPR050524) [family] |
| *CPAR2_600940* | WT (145) <sup>1</sup><br>**c.859-911del (226)** <sup>3</sup><br>c.828del (1) <sup>3</sup> | N/A | Bul1-like (IPR039634) [family] |
| *CPAR2_800150* | WT (51) <sup>1</sup><br>**c.298-299del (313)** <sup>3</sup><br>C57A, Y19* (7) <sup>2</sup><br>c.292del2, c.298-299del (1) <sup>3,4</sup> | *AQY1* | Aquaporin transporter (IPR034294) [family] |

<sup>1</sup> Wildtype allele.
<sup>2</sup> Non-synonymous substitution. The nucleotide difference from the wildtype sequence is written before the arrow, and the amino acid change is written after the arrow.
<sup>3</sup> Deletion in coding sequence, at the nucleotide positions indicated, resulting in a frameshift.
<sup>4</sup> This allele contains separate 1-bp and 2-bp deletions near each other. The net effect is that the open reading frame is maintained and there is no premature stop codon.

*DAL4* (CPAR2_103200) is predicted to encode an allantoin permease due to its similarity to *S. cerevisiae DAL4*. The Q109* nonsense mutation is homozygous and is found only in isolates from one branch of Clade 4, which includes the reference strain CDC317 (Figure 1) (37, 38). Owing to this, CPAR2_103200 is classified as a pseudogene in the Candida Genome Database (39).

CPAR2_401360 (*AGP2*) encodes a proline permease of 575 amino acids, unlike the allele (428 amino acids) in CDC317 and other Clade 4 isolates which results from a deletion of nucleotides 425-426 (Figure 1). Separately, the isolates NRZ-BK2179 and NRZ-BK982 also encode a truncated Agp2 protein resulting from a stop codon gain (Y117*) in the gene (Figure 1, Table 1).

CPAR2_600940 is a member of a Bul1-like family. In *S. cerevisiae*, Bul1 is a subunit of the ubiquitin ligase complex (36). Short forms of CPAR2_600940 are caused by deletion of nucleotides 859-911 in isolates in Clades 3, 4, and 5, as well as isolates NRZ-BK2179 and NRZ-BK982, and separately by deletion of nucleotide 828 in one isolate (NRZ-BK680) from Clade 3 (Figure 1, Table 1).

CPAR2_800150 is orthologous to the aquaporin gene *AQY1* of *C. albicans* (31) (Table 1). *AQY1* is truncated in the majority of the 372 isolates by a deletion of nucleotides 298-299 (Figure 1, Table 1). The distribution of this variant implies it occurred independently twice; once in the ancestor of Clades 4 and 5, and once in the ancestor of Clade 1 (Figure 1). In NRZ-BK1935, this frameshift is compensated by a further deletion of nucleotide 292, restoring the reading frame (Figure 1, Table 1). The longer, wild-type *AQY1* allele is found in all isolates of Clade 3, and in many isolates in Clade 2 (Figure 1, Table 1). Isolates on one branch of Clade 2 have truncated *AQY1* alleles resulting from a stop-gain variant (Y19*) (Figure 1, Table 1).

In these four cases, the effect of the internal stop codon or frameshift was so severe that the two forms fell into two different sequence clusters. This suggested that stop codon variants may exist in other genes too. We identified homozygous stop gain-variants in 183 genes that altered protein length but did not impact gene clustering (Supplementary Table S1D).

### Expansion/contraction of tandem arrays

The accessory genome is enriched for members of the Major Facilitator Superfamily (MFS) of transporters (Supplementary Table S1C, Supplementary Table S1E). There are 24 MFS genes in the accessory genome, of which eight are homologs of the inorganic sulphur transporter *SOA1*. A locus containing a tandem array of five *SOA1* transporters was previously annotated in the reference genome (CPAR2_702920, CPAR2_702940, CPAR2_702950, CPAR2_702960 and CPAR2_702970) (Figure 4A). Our analysis now suggests that the reference gene CPAR2_702960 results from a fusion of two paralogs (which we named PARALOG_702960_1 and PARALOG_702960_3) that occurred in an ancestor of the reference isolate CDC317 and 8 other isolates (Type 2A in Figure 4). Two different fusions between PARALOG_702960_1 and PARALOG_702960_3 were identified that differ at the fusion point (Supplementary Figure 4). One fusion, Type 2A, is the structure seen in the reference strain and others in Clade 4. The other fusion, Type 2B, is seen in 29 isolates in Clade 3 (Figure 4). These fusion events likely occurred independently, and they both resulted in the deletion of a formate dehydrogenase (*FDH*) gene that lies between the paralogs. Most isolates (334) have 6 copies of *SOA1* at this locus, and this arrangement likely represents the ancestral state of the locus (Type 1 in Figure 4A). We identified three other *SOA1* array structures that we named array Types 3, 4, and 5, occurring in two isolates each (Figure 4). Each of these array structures involves fusions between different *SOA1* genes in the array.

**Figure 4.**
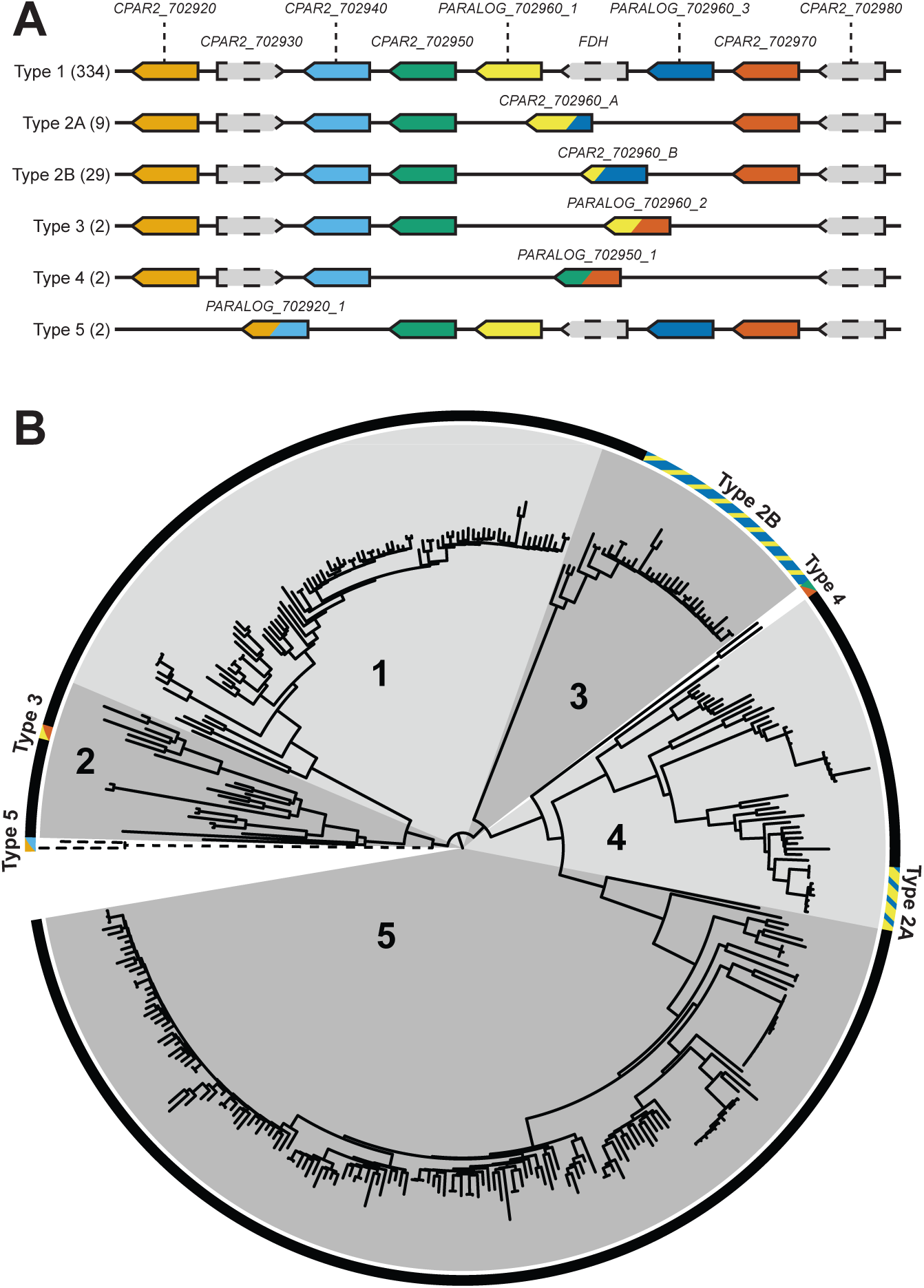
Organisation of the SOA1 locus in the *C. parapsilosis* pangenome. (A) Five tracks show the differing gene content at the *SOA1* locus caused by fusion rearrangements. *SOA1* paralogs are in colour, and fusions between paralogs contain the colours of both original genes. Unrelated genes are shown in grey with dotted borders. Type 1 represents the likely ancestral state. Fusions between PARALOG_702960_1 and PARALOG_702960_3 occurred in isolates in Type 2A and Type 2B, between PARAL-OG_702960_1 and CPAR2_702970 in isolates in Type 3, between CPAR2_702950 and CPAR2_702970 in Type 4, and between CPAR2_702920 and CPAR2_702940 in Type 5. The number of isolates with each Type organisation is shown in brackets. (B) The tree in Figure 1 is replicated here, with additional annotation showing the SOA1 locus type for each strain.

The variation in copy number of *SOA1* genes suggests that *C. parapsilosis* isolates may vary in their ability to transport inorganic sulphur. Analysis of RNA-seq data from (40) shows that expression of PARALOG_702960_1, PARALOG_702960_3 and CPAR2_702970 is significantly upregulated by sulphur limitation (absence of cysteine and methionine in media) (p ≤ 2.58E-05) with log2 fold changes ranging from ∼2.6 to ∼7 (Supplementary Table S2). Expression of other *SOA1* paralogs at this locus is not affected (Supplementary Table S2).

Variation in *ALS* (adhesin) gene content resulting from gene fusion in *C. parapsilosis* has been described previously (15). All five known *ALS* genes (*ALS1, ALS3, ALS6, ALS7,* and *ALS11*) are present in 369 *C. parapsilosis* isolates. We found a fusion between *ALS11* and *ALS7* (PARALOG_404800_1) in one isolate in Clade 2 (MSK2386) and two isolates in Clade 5 (MSK809 and FM09), that also resulted in the deletion of *ALS6*. A similar fusion was described by Pryszcz et al. (15), suggesting that similar fusions have occurred multiple times.

We observed similar fusions between tandemly duplicated genes, involving homologs of *JEN1* and of *SIT1* (both members of the MFS family), and in a several short-chain dehydrogenase genes (*AYR, SCR2, SCR3,* and *DLD1*) (Supplementary Table S1C). We found other reference genes that likely result from fusions between tandem paralogs including *JEN2* (another MFS gene), *PLB3*, and *SNG1*. The majority of the differences in gene content between *C. parapsilosis* strains result from the expansion or retraction of these tandem gene arrays.

### Association between accessory genes and phenotypes

To determine whether there are associations between the presence/absence of accessory genes and phenotype, we looked for correlations of genotype with growth on 23 nitrogen sources, using the program TreeWas (41). For testing nitrogen sources, phenotypes were scored as log2 ratios of growth on the control medium (0.5% ammonium sulphate) compared to growth on 10 mM concentration of 23 other sole nitrogen sources (Supplementary Table S3). Only growth in two conditions (lysine and proline) showed significant associations (Supplementary Table S4).

The presence of fusion gene CPAR2_701140, and its parental sequences PARALOG_701140_1 and PARALOG_701140_2, were strongly associated with growth on proline as a sole nitrogen source (p = 0) (Supplementary Table S4). These are orthologs of *S. cerevisiae* phospholipase *PLB1*, deletions of which have been associated with a reduced capacity to utilise proline as a nitrogen source (42). Improved growth using proline as a sole nitrogen source is correlated with presence of the fusion gene CPAR2_701140, i.e. with reduction from two copies to one copy by fusion (Supplementary Table S4). The gene fusion is observed in all isolates in Clades 4 and 5, and in two branches of Clade 2. Among the 372 *C. parapsilosis* isolates, 176 have log2 growth ratios ≥1 on proline, i.e. growth is double or better relative to control, and 143 of these have the fusion gene CPAR2_701140. The majority of these isolates are found in Clade 5.

The absence of three genes (CPAR2_204210, CPAR2_807730, CPAR2_807740), and the presence of three other genes (PARALOG_807730_1B, PARALOG_402040_1, PARALOG_402040_2) were strongly correlated with reduced growth on lysine as a sole nitrogen source (p = 0) (Supplementary Table S4). None of these genes nor their *S. cerevisiae* orthologs are known to have functions in lysine utilisation. CPAR2_204210 is a homolog of *C. albicans* alkane-inducible cytochrome P450 gene *ALK8*. PARALOG_807730_1B results from the fusion of CPAR2_807730 and CPAR2_807740 (i.e. reduction from two genes to one), and this fusions occurs in Clades 2, 4, and 5. They are homologs of the *S. cerevisiae* D-lactate dehydrogenase gene *DLD1.* Null alleles of *S. cerevisiae DLD1* result in reduced capacity to utilise arginine as a nitrogen source (42), but a phenotype involving lysine utilization has not been reported in *S. cerevisiae*. PARALOG_402040_1 and PARALOG_402040_2 are orthologs of *C. albicans* dicarboxylic acid transporter gene *JEN2*. Among the 372 *C. parapsilosis* isolates, 61 have log2 ratios ≤ −1 on lysine (i.e. growth is halved or worse relative to control). The majority of these isolates are found in Clade 5. Accessory genes CPAR2_204210, CPAR2_807730, and CPAR2_807740 are all present in only five of these 61 isolates while genes PARALOG_807730_1B, PARALOG_402040_1, and PARALOG_402040_2 are found in 56, 61 and 54 isolates respectively.

To test whether strains with similar phenotypes and similar gene content are also more closely related to each other, we measured the phylogenetic signal of these genes using Blomberg’s K-values, treating presence or absence of an accessory gene as a binary trait. The phylogenetic signal for the presence of these genes is high (Table 2), suggesting that fusions, gene gains and gene losses occurred in ancestors of clades and were inherited by their descendant isolates. Because these genotypes are very dependent on phylogeny it is difficult to assess if their correlations with phenotype is real or confounded by shared ancestry. While several associations between gene content and phenotype reached significance in this GWAS, it is difficult to be certain whether these differences in gene content contribute to phenotype or have inflated association scores owing to the clonal population structure of *C. parapsilosis*.

**Table 2:**
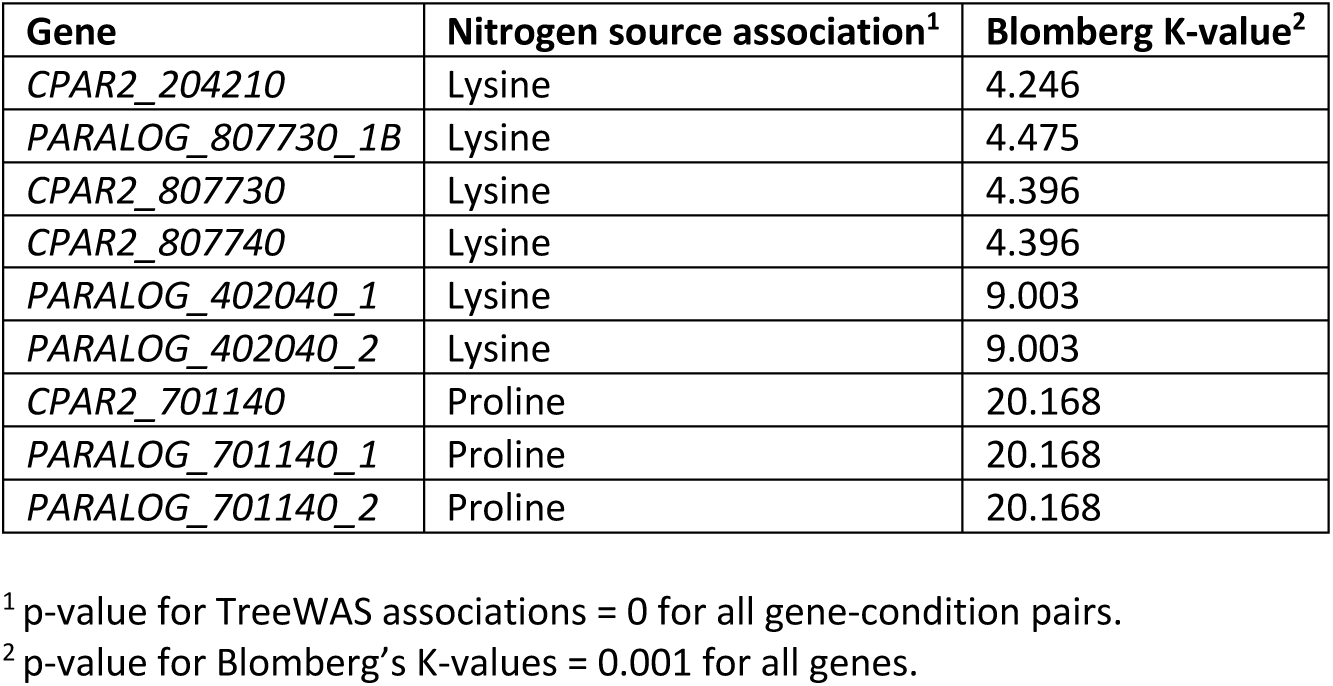
Genes significantly associated with growth on specific nitrogen sources.

| Gene | Nitrogen source association<sup>1</sup> | Blomberg K-value<sup>2</sup> |
| --- | --- | --- |
| *CPAR2_204210* | Lysine | 4.246 |
| *PARALOG_807730_1B* | Lysine | 4.475 |
| *CPAR2_807730* | Lysine | 4.396 |
| *CPAR2_807740* | Lysine | 4.396 |
| *PARALOG_402040_1* | Lysine | 9.003 |
| *PARALOG_402040_2* | Lysine | 9.003 |
| *CPAR2_701140* | Proline | 20.168 |
| *PARALOG_701140_1* | Proline | 20.168 |
| *PARALOG_701140_2* | Proline | 20.168 |

<sup>1</sup> p-value for TreeWAS associations = 0 for all gene-condition pairs.
<sup>2</sup> p-value for Blomberg's K-values = 0.001 for all genes.

### *DAL4* (CPAR2_103200) truncation impacts allantoin metabolism

*DAL4* truncation failed to show association with the phenotypes tested in the GWAS analysis. However, these results may have been confounded by strict population structure that might have obscured actual association. In *S. cerevisiae*, null alleles of *DAL4* result in a failure to uptake allantoin and use it as a nitrogen source (43). To test whether the truncated allele of *DAL4* in *C. parapsilosis* correlates with the inability to use allantoin, we compared the growth of Clade 4 isolates with homozygous Q109^WT^ and Q109* alleles on media containing allantoin as a sole nitrogen source, relative to media containing ammonium sulphate (Figure 5). All Clade 4 isolates preferentially used allantoin rather than ammonium sulphate (positive log2 ratio). However, the log2 ratios were significantly lower in alleles containing the *DAL4* Q109* (Figure 5A).

**Figure 5.**
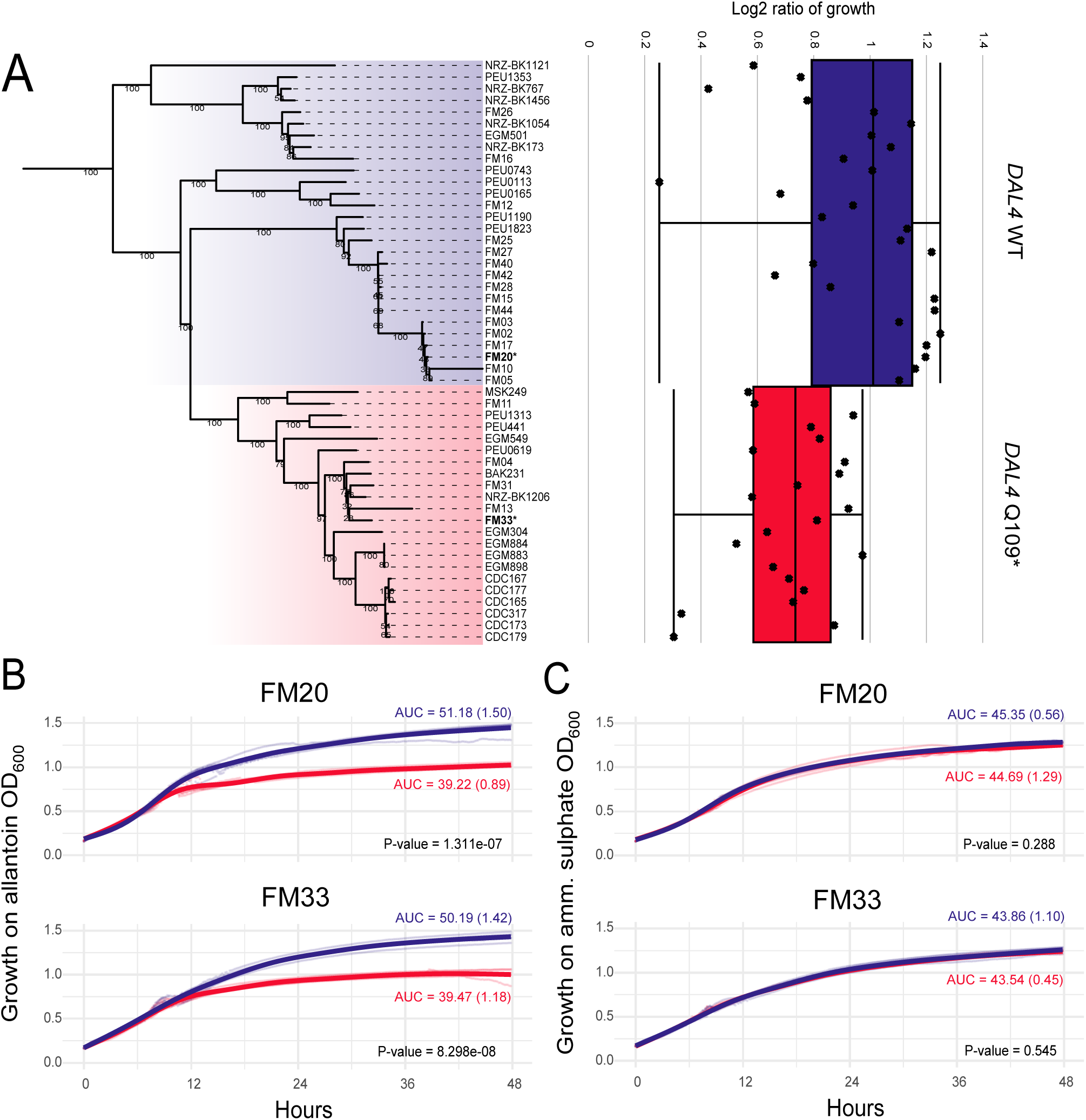
Clade 4 isolates with truncated DAL4 show reduced growth on allantoin. (A) Left: The phylogeny of Clade 4 is shown with all other clades collapsed. All isolates containing the DAL4 Q109* allele are highlighted in red. Other Clade 4 strains are highlighted in blue. Right: Boxplots showing growth of Clade 4 isolates on allantoin grouped by DAL4 genotype. Boxes display the inter-quartile ranges, and whiskers show the range of outliers. Individual isolalte values are jittered with respect to their position on the phylogenetic tree to the left. (B) Top: Comparison of growth on allantoin of *C. parapsilosis* FM20 (WT) in blue, with the Q109* variant added using CRISPR-Cas9 (red). Bottom: Comparison of growth on allantoin of *C. parapsilosis* FM33 (Q109*) in blue, with the Q109* variant removed using CRIS-PR-Cas9 (red). Mean values for area under curve (AUC) are shown, with standard deviation in parentheses for each curve. P-values for differences between AUC values. (C) Same as panel B, with growth on ammonium sulphate.

To test if the Q109* variant in *DAL4* directly affects function, CRISPR-Cas9 gene editing was used to introduce the stop codon into *DAL4* in *C. parapsilosis* FM20 (which is homozygous for the wild-type allele), and to remove the stop codon in *C. parapsilosis* FM33 (which is homozygous for the Q109* allele). Addition of the early stop codon in the FM20 background resulted in reduced growth on allantoin as a sole nitrogen source (Figure 5B), while removal of the stop-gain in the FM33 background resulted in improved growth on allantoin as a sole nitrogen source (Figure 5B). Neither edit affected growth in ammonium sulphate (Figure 5C).

Although truncating *DAL4* in *C. parapsilosis* results in reduced utilisation of allantoin, growth is not completely abolished, unlike *DAL4* null mutants of *S. cerevisiae* (43). Because *DAL4* is only truncated and not deleted, this may explain a less severe phenotype. Further investigation of InterPro (44) results identified three other putative allantoin permease genes: CPAR2_201790 and CPAR2_201900, both homologous to *FCY21*, and CPAR2_806580, orthologous to *FCY23* in *C. albicans.* These additional transporters may explain a lack of strong phenotype associated with loss of function of *DAL4*, although orthologs of *FCY21* are also present in *S. cerevisiae* (45).

### Gene content of divergent strains does not contribute highly to pangenome

The divergent strains PEU651 and EGM1079 were excluded from systematic pangenome analysis to avoid biases introduced by outliers. Instead, comparisons of gene content in these strains were made separately, using two approaches. Firstly, BLASTN was used to identify genes annotated in PEU651 and EGM1079 that match the 5859 gene clusters from the other isolates. Secondly, to identify genes that were incorrectly called as absent in the divergent strains due to annotation errors, short reads from PEU651, EGM1079, and CDC317 were aligned to the best respective assemblies of the other two strains, after which coverage analysis was performed to check whether genes in each assembly were present or absent in the other strains.

We found only minor differences between the gene content of *C. parapsilosis* PEU651 and EGM1079 and the other 372 isolates (Supplementary Table S5). Of the 26 hypothetical novel genes found in the 372-isolate pangenome analysis, eight are present in PEU651 and 11 are present in EGM1079. Several gene fusions were identified including one at the *SOA1* locus resulting in novel array Type 5 (Figure 4). A total of 17 genes present in the CDC317 reference are predicted to be absent in PEU651 and EGM1079 based on coverage, including three blocks of contiguous genes. Four genes annotated in PEU651, and six genes annotated in EGM1079, are predicted absent in CDC317 by coverage analysis (Supplementary Table S5).

### Three unique inversions in the genomes of divergent strains

Genomic rearrangements between the two divergent strains and the CDC317 reference were investigated by aligning their long-read assemblies to the reference, resulting in the discovery of three inversions, all between 30 and 40 kb in size and flanked by inverted repeats (Figure 6).

**Figure 6.**
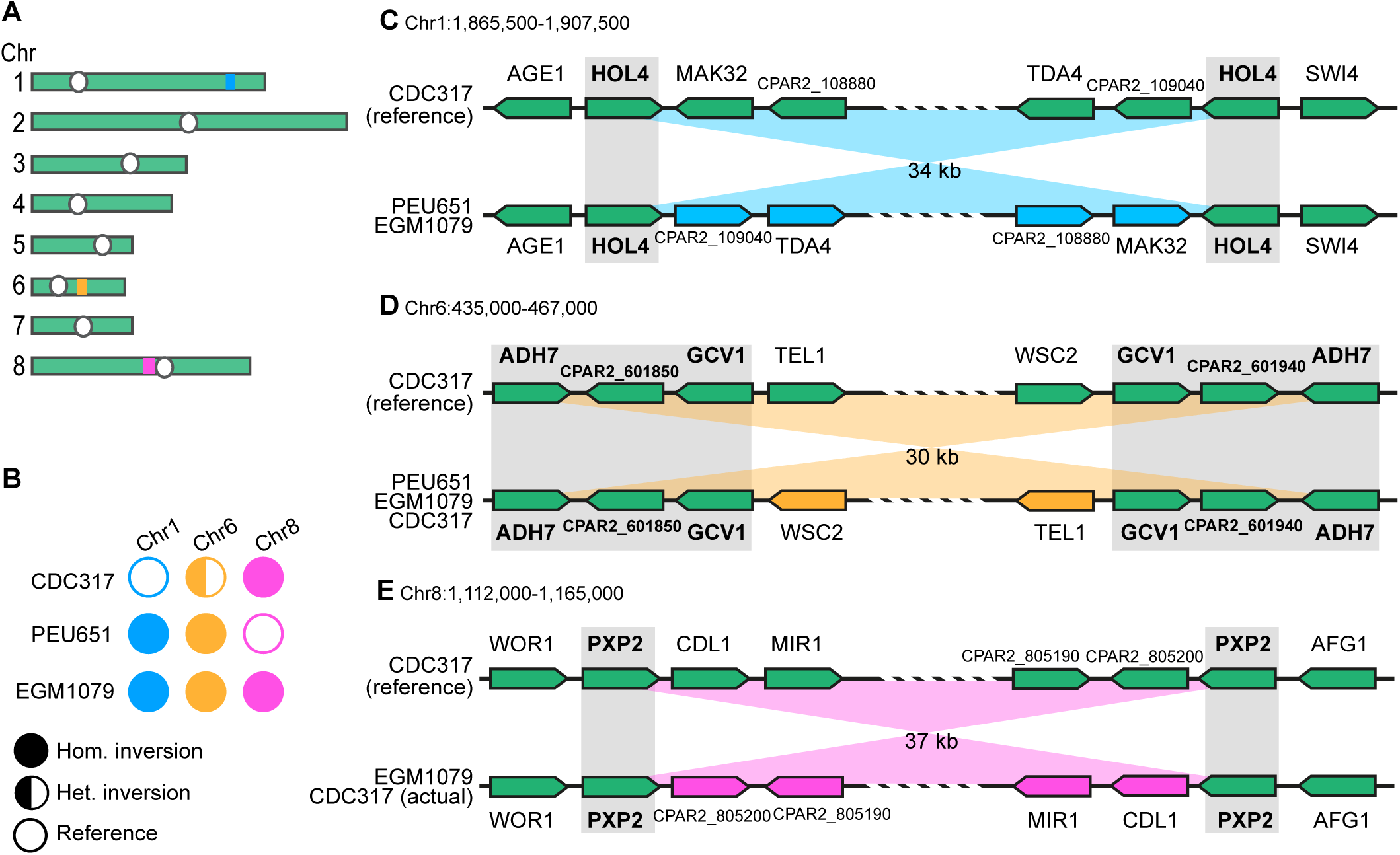
Inversions in the divergent *C. parapsilosis* isolates EGM1079 and PEU651. (A) Locations of the three inversion regions on chromosomes 1 (blue), 6 (orange) and 8 (magenta). White circles indicate the centromeres. (B) Inference of homozygosity or heterozygosity of the inversions in different isolates, based on analysis of MinION long reads spanning the endpoints of each inverted region. Fully colored circles indicate that all the reads support the presence of the inversion. Half-colored circles indicate that approximately half the reads support the presence of inversion but the other half do not, i.e. that the strain is heterozygous for the two orientations of the region. Empty circles indicate that no reads support the presence of the inversion. (C-E) Gene organization at the endpoints of the inversion regions on (C) chromosome 1, (D) chromosome 6, and (E) chromosome 8. Gray shading and bold gene names indicate the regions that form inverted repeats (>99.5% sequence identity) on each chromosome.

First, PEU651 and EGM1079 both have a homozygous 34-kb inversion on chromosome 1 compared to the reference, with breakpoints in a pair of identical copies of the gene *HOL4,* which form an inverted repeat (Figure 6C).

Second, PEU651 and EGM1079 both have a homozygous 30-kb inversion on chromosome 6, with breakpoints in an inverted repeat formed by two near-identical (99.5%) copies of a 3.3-kb sequence containing three genes: *ADH7*, an uncharacterised gene, and *GCV1* (Figure 6D). By aligning MinION reads from *C. parapsilosis* CDC317 (47) to the reference sequence, we found that this 30-kb inversion is heterozygous in strain CDC317, but the haploid reference genome sequence derived from it contains the orientation allele that is absent from the two divergent strains (Figure 6B).

Third, when we mapped MinION reads from *C. parapsilosis* CDC317 (47) to the reference genome sequence we found that all the reads support the presence of a 37-kb inversion on chromosome 8 (Figure 6B,E) relative to the reference, which suggests that there is an assembly error in the reference. Remarkably, however, the orientation of this 37-kb region differs between the two divergent *C. parapsilosis* isolates EGM1079 and PEU651, so its orientation is polymorphic in the *C. parapsilosis* population. The endpoints of this inversion are in two near-identical copies of the gene *PXP2*. A possible explanation for this situation is that the culture of *C. parapsilosis* CDC317 from which the reference genome sequence was obtained in 2009 by Sanger sequencing may have been heterozygous for the two orientations of this 37-kb region, resulting in one orientation being chosen for use as the reference, but the culture of *C. parapsilosis* CDC317 from which the MinION reads were obtained (2022, (47)) may have undergone loss of heterozygosity in the region and retained only the allele whose orientation is opposite to the reference.

Because these three inversions do not alter the gene content of *C. parapsilosis* EGM1079 and PEU651, and their breakpoints occur in near-identical genes whose coding frames remain intact, it is unlikely that they have a major effect on phenotype.

## Discussion

The *C. parapsilosis* pangenome is considerably smaller than that reported for other yeasts and fungi. Our sample set of 372 genomes is similar in size to the one that was used to define the pangenome of *Aspergillus fumigatus* (300 genomes) (20). However, the pangenome of *C. parapsilosis* is much smaller; we identified 5,859 homologous genes clusters of which 98.8% are core genes compared to 10,907 gene clusters in *A. fumigatus*, of which 69% are core genes (20). The pangenome of *C. albicans* was estimated to consist of 7,325 gene clusters (∼90% core), and the pangenome of *Cryptococcus neoformans* is 8,193 gene clusters (∼80% core) (18). For *S. cerevisiae*, estimates of gene clusters in the pangenome range from 7,708 to 7,796, with 63-85% of these being core genes (18, 24, 25). The variation in the *S. cerevisiae* calculations depend on the number of genomes included and methodology used (18, 24).

Whereas the average number of gene models per strain in *C. parapsilosis* (5,849) is similar to that of *C. albicans* (5,993) and *S. cerevisiae* (5,759) (18), the accessory component in *C. parapsilos*is is much smaller (Supplementary Figure 5). The major outstanding question is whether this difference in the estimated size of the accessory genome reflects underlying biological differences, or differences in the methods and definitions that were used to quantify the pangenome in each species. *Candida glabrata* was found to have a small accessory component with 6% (323 genes), which the researchers found shrunk following manual curation of fragmented assemblies (46). In the *C. albicans* pangenome, 1013 duplicates of core genes were identified (18) which may be equivalent to the 74 mixed clusters we describe in *C. parapsilosis*. We assigned the 152 as core genes, whereas similar clusters in *C. albicans* may have been counted as accessory genes. The *C. albicans* accessory genome may therefore be inflated due to differences in terminology used.

However, the accessory genome of *C. parapsilosis* is still surprisingly small (1.2 %, 68 genes), which reflects the general lack of sequence diversity (0.16 to 0.95 heterozygous sites compared to 3.2 to 9.9 per kb in *C. albicans* (8)). The clonal nature of *C. parapsilosis* is likely amplified by dysfunction of its sexual cycle; the 374 *C. parapsilosis genomes* characterized here contain only the *MTL*a idiomorph.

Despite the small size of the accessory genome in *C. parapsilosis*, we identified expansion and contraction of MFS genes, which also occurs in the accessory genomes of *C. albicans* and *A. fumigatus* (18, 20). We also observed expansion and contraction of FAD binding domain proteins in the accessory genome, similar to *A. fumigatus* (Supplementary Table S1C) (20). These families may therefore be subjected to selection pressure in several species.

We found that duplication and gene fusion are common mechanisms of gene gain and loss in *C. parapsilosis*, including multiple independent fusions between *SOA1* paralogs. We previously described similar fusions of tandem copies of the ABC drug transporter *CDR1B* that likely contribute to antifungal resistance (47). In addition, Pryszcz et al. (15) described four different recombination events at *ALS* loci, three involving different fusions between *ALS3*, *ALS11* and *ALS7*, and another between *ALS1* and *ALS3*. In our analysis, we identified three separate instances of fusion between *ALS11* and *ALS7*. Frequent independent tandem amplification of *RTA3* has also been demonstrated (14). Our analysis supports the hypothesis that gene duplication and gene fusion is a dominant feature in the evolution of metabolic diversity in fungi (48). We also suggest that tandem amplification and subsequent fusion is a main driver of the accessory genome.

We identified several issues resulting from using a single reference genome sequence, i.e. the *C. parapsilosis* CDC317 reference assembly, which was originally generated using Sanger sequencing (1). In this reference sequence, *DAL4* and three other genes were classified as pseudogenes because of the presence of a premature stop codon (1), but they are intact in other isolates. The presence of truncated *DAL4* alleles in several related clinical strains may indicate a lack of pressure for allantoin use in *C. parapsilosis*, at least in clinical settings. We also identified premature stop codons in 183 other genes in other isolates (Supplementary Table S1D). The CDC317 reference does not encode any of the 30 novel genes we identified, whether paralogs of annotated genes recently duplicated, or unannotated ORFs. Studies performed by comparison to only the reference genome may miss variants in these genes.

Of the 23 nitrogen metabolism phenotypes we investigated using TreeWAS, only two showed significant correlations (41). All the associated genes had very high Blomberg’s K values, suggesting that the presence or absence of the gene is shared across many closely related isolates. This pattern implies that any phenotype strongly associated with the presence of one of these genes would, in fact, also be strongly associated with every genomic characteristic shared by these closely related isolates, making it difficult to disentangle causation from correlation. The lack of variation in the *C. parapsilosis* pangenome also reduces the power of association studies; only 68 high-quality gene presence/absence polymorphisms were identified.

We also discovered a pair of distantly related *C. parapsilosis* isolates, EGM1079 and PEU651. These isolates are still ∼99.6% identical to the other 372 isolates, which is similar to the variation between strains of other yeast species (49). Comparisons between orthologs in the *C. parapsilosis* species complex shows a typical nucleotide identity of 82-86% (50). The strains are unlikely to be hybrids, as has been found elsewhere in the *C. parapsilosis* species complex (51, 52), because the heterozygosity levels are low. The vast majority of *C. parapsilosis* isolates sequenced to date from clinics all over the world belong to a single lineage with five related clades. However, we have found two strains belonging to a separate lineage, EGM1079 and PEU651, suggesting that increased global sampling of *C. parapsilosis* isolates may yet identify further rare human-associated populations. Understanding the distribution of these lineages could provide deeper insights into the virulence and epidemiology of *C. parapsilosis*, especially in diverse clinical settings.

## Methods

### Genome sequencing and data acquisition

The genomes of 374 *C. parapsilosis* isolates were used in this analysis, including some from previously published genomes (Supplementary Table S1A). Illumina short-read sequencing of genomes reported in this study were sequenced as described in Bergin et al. (14). In brief, total genomic DNA was extracted using phenol-chloroform-isoamyl alcohol. Library preparation and sequencing was performed in 150 bp paired-end format using the corresponding library preparation kits and Illumina instruments as described in Supplementary Table S1A.

For Nanopore sequencing, high molecular weight DNA was extracted using the Biosearch Technology Masterpure Yeast DNA purification Kit (MPY80010). Sequencing was performed with 1 µg of DNA using the Native Barcoding Kit (SQK-NBD114-24) (Supplementary Table S1A).

### Raw read processing, assembly and filtering

Short-reads were trimmed and filtered using Skewer v. 0.2.2 (53) to minimal mean qualities of 30 and minimal lengths of 35. Assembly of short-reads was performed using SPAdes v. 3.14.0 (27) using default settings. Assemblies containing scaffolded contigs were used. Scaffolds < 500 bases or of quality < 10 were removed. For Nanopore reads, filtering was performed using NanoFilt v. 2.8.0 (ONT) to remove reads with Q < 7 and L < 1kb. Nanopore assemblies were generated using Canu v. 2.2 (28). Contig polishing was performed using Illumina short-reads with NextPolish v. 1.4.1 (54). Five rounds of correction were performed. Contigs were compared to the *C. parapsilosis* CDC317 reference genome using dot-plot-matrices in D-Genies (55). Manual joins were made in some contigs.

Contigs were assigned to chromosomes and re-orientated with respect to the *C. parapsilosis* CDC317 reference genome using BLAST v. 2.10.0+ (56) and BedTools v. 2.30.0 (57). Contigs that could not be assigned to the reference were manually investigated to assess if they represented novel sequences or likely contamination. Novel sequences were maintained while contaminants (i.e. sequences whose likely origin was not yeast or bacteria according to NCBI BLAST (58), or showed no evidence of integration) were removed. All mitochondrial contigs were removed. Softmasking of repetitive regions was performed prior to gene annotation using both RepeatModeler v. 2.0.4 and RepeatMasker v. 4.4.2-p1 (59).

### Gene annotation and ortholog clustering

Each genome was annotated using BRAKER3 v. 3.0.2 (30). The training set for protein annotation was derived from the Candida Gene Order Browser (31) using protein clusters present in three or more species. The parameters used with BRAKER3 were “-- translation_table=12”, “--fungus”, and “--downsampling_lambda=0” to facilitate annotation characteristics of CUG-Ser1 fungal species that have few introns. Nucleotide gene sequences were grouped into homologous clusters using GET_HOMOLOGS v. 07122022 (17) with default settings.

### Assignment of homologous clusters to *C. parapsilosis* CDC317 reference genes (Supplementary Figure 1 and 2)

The nucleotide sequences of each homologous cluster were translated and aligned using MAFFT v. 7.520 (60) with the following parameters: “--maxiterate 1000 --op 1.0 – genafpair”. Hidden markov models (HMMs) of each cluster were generated using HMMBuild v. 3.3.1 (61). HMMs for each cluster were assigned to reference proteins from *C. parapsilosis* CDC317 using Orthofisher v. 1.0.3 (62). Homologous clusters were assigned based on the highest scoring reference protein. Clusters with equal scores to multiple reference proteins, clusters which shared the highest score for the same reference protein, or clusters lacking scores for any reference protein were manually investigated. Gene models within unique clusters (clusters composed of sequences derived from only one or two genomes) were manually investigated using BLASTN (56) to compare to the *C. parapsilosis* CDC317 reference gene set. If a significant hit could be found to a reference gene, the unique cluster was combined with the cluster representing that gene. Those not represented in the reference set were manually investigated. Clusters consisting of short proteins (<100 amino acids) with no assignment to the CDC317 reference gene set, or which did not contain any functional domains identifiable by InterProScan (44) were removed.

In some cases, two or more homologous clusters represented the same gene sequence but were annotated differently by BRAKER3. This was because of the presence of premature stop codons in the open reading frames in some genomes. BRAKER3 often deals with stop gains by inserting artificial introns, or by splitting the sequences on either side of the stop codon to generate two genes. The variation in sequence output often splits what should be one homologous cluster into at least two separate clusters.

Homologous clusters were merged and split where appropriate to fix misannotations, distinguish paralogues and fusion genes, and identify paired clusters with and without premature stop codons where appropriate. Not all clusters could be resolved to likely single orthologous sequences. Due to limitations in short-read sequencing, assembly tends to fail at identical or near-identical sequences at multiple loci often generating multiple fragments of protein sequences. Fragments are often clustered together by GET_HOMOLOGS, and it can be difficult-to-impossible to distinguish which locus each fragment derives from. Presence and absence of unresolved orthologues was determined by comparing read coverage at each locus using short-read coverage against either the CDC317, NRZ-BK680or CLIB214 reference genome. Copy number of all representative loci per homologous cluster were determined using Delly cnv v. 0.8.7 (63) and BedTools intersect (57) using BAM files generated for SNP calling. Presence of the locus was assumed when copy number was at least one.

### Gap filling of homologous clusters using BLAST

Preliminary analysis of the presence or absence of short gene sequences (homologous clusters with the sequences <500 bp) revealed an often-random pattern across the 372 genomes. This often resulted from gaps in the annotation, for example if the genes lay at the ends of assembled contigs. To fill arbitrary gaps in annotation, presence/absence assignments were further refined through BLAST (56) analysis using representative sequences for each group. Representative sequences were chosen from the CDC317 reference genome where available, or from one of seven other chromosome-level assemblies (MSK478, MSK802, MSK803, MSK812, CLIB214, NRZ-BK680 or UCD321) if not. If no such sequence was available, the longest nucleotide sequence for that homologous cluster that contained no sites of scaffolding (no Ns in sequence) was selected. Presence in a genome was assigned based on 99% identity and 99% match length. This was applied to all homologous clusters resolved to likely single orthologues.

### Confirming presence/absence of homologous clusters using expected gene order

To further confirm the absence of specific homologous clusters in individual isolates, the presence of genes on either side of the target was determined. Gene order of all resolved homologous clusters (*C. parapsilosis* CDC317 reference genes and novel sequences) was determined in each genome to assign a consensus gene order across all isolates. Gene absence in a specific genome was assumed if the expected neighbouring genes were present. The shortest gap in annotation between two expected neighbouring genes on the same contig was identified. If this was not possible, or if the sequence between the neighbouring loci contained a series of Ns indicating a point of scaffolding, the gene was not assumed to be absent. Genuine absence was defined by the presence of neighbouring genes with an intact sequence between them. All homologous clusters resolved to likely single orthologues were tested in this way. Presence and absence of neighbouring genes within the same gene family were manually investigated to assess potential gene fusion events. The accessory genome is defined as genes absent in more than one genome. Blomberg’s K-values were determined for each accessory gene in R-package PhyTools v.2.1-1 (35) using the phylosig function. P-values were determined under 1000 permutations.

### SNP calling and variant analysis

Short reads were mapped to reference genomes CDC317, CLIB214 and NRZ-BK680 using BWA mem (64). Alignments were sorted and indexed using SAMTools (65). Duplicate reads were marked using Picard tools (66). BAM files for CLIB214 and NRZ-BK680 were used in coverage analysis only, while the BAM file for CDC317 was also used in SNP calling. Variants were called in parallel using GATK (v. 4.0.1.2) (66) module HaplotypeCaller, freebayes (v1.3.9) (67), and bcftools mpileup (v1.10.2) (65). Results were merged using bcftools isec to keep only variants called by two or more of the tools. Variants were filtered using GATK VariantFiltration (66) using the following parameters: minimum mapping qualities of 30, minimum genotype qualities of 40, minimum read depth of 15. Additionally, only biallelic SNPs were retained. The predicted effects of variants found not present in the CDC317 reference genome were analysed using a SIFT database generated for the *C. parapsilosis* CDC317 reference genome (14, 68).

### Phylogenetic analysis

A FASTA alignment of all sites containing an SNP in at least one isolate was created from the multi-sample VCF file using a custom script (https://github.com/CMOTsean/HetSiteRando). Heterozygous variants were randomly assigned to either allele on a per-site basis. An SNP tree was constructed with the alignment file using RAxML version v8.2.12 with the GTRGAMMA model of nucleotide substitution and 1,000 bootstrap replicants (26).

### Functional analysis using InterProScan

InterProScan v. 5.61-93.0 (44) was used to further annotate putative protein function using PANTHER, TIGRFAM, PFAM and SUPERFAMILY databases for all representative sequences. GO analysis of the accessory and total gene sets was determined by hypergeometric test for overrepresentation using SciPy (69).

### Phenotypic analysis

*C. parapsilosis* strains were grown from stock on YPD agar (1% Bacto Yeast Extract (212750, Sigma), 2% Bacto Peptone (211677, Sigma), 2% Bacto Agar (214010, Sigma), 2% D-(+)-Glucose (G8270, Sigma)) and incubated at 30°C for 48 h. Biological replicates from each strain were inoculated in 150 µl YPD broth (1% Bacto Yeast Extract (212750, Sigma), 2% Bacto Peptone (211677, Sigma), 2% D-(+)-Glucose (G8270, Sigma)) in flat-bottom 96 well plates. Type strain *C. parapsilosis* CLIB214 was inoculated into the bordering columns and rows of each plate, around the first two and last two columns and rows, to reduce edge effects. These replicates were ignored in in downstream analyses. In total, eight plates were used. Plates were incubated at 30°C, 1100 rpm for 24 h. Stock plates were made by mixing 70 µl of each culture was with 70 µl 30% glycerol, which were then stored at −80°C. To test strains in different conditions, stock plates were defrosted and pinned onto YPD agar in PlusPlates using a ROTOR HDA robot and incubated at 30⁰C for 24 h. Colonies were inoculated into 150 µl YPD broth in 96 well cell culture plates using the robot and incubated at 30°C, 1100 rpm for 24 h. Culture plates were combined in duplicate in a 1536 format by pinning to YPD agar on PlusPlates using the ROTOR HDA robot and incubated at 30°C for 24 h. Colonies were then pinned in tetrads: sets of 4 replicates (two biological replicates and two technical replicates) to control plates (0.19% yeast nitrogen base with no amino acids or ammonium sulphate, 0.5% ammonium sulphate, 2% glucose, 2% agar) and 47 different test plates (0.19% yeast nitrogen base with no amino acids or ammonium sulphate, 2% glucose, 2% agar, and 10mM of a single nitrogen source for tested nitrogen sources, or 0.5% ammonium sulphate for testing other compounds). Other tested compounds and concentrations are listed in Supplementary Table S4. Plates were incubated at 30°C and photographed after 48 and 72 hours of incubation. A Singer Phenobooth was used to count the number of pixels per replicate in each tetrad. Median values per tetrad were calculated and normalised per plate by subtracting the plate-wide median. The log2 ratios of each strain’s median values between test and control plates were used as growth values. Z-scores of log2 ratios were calculated on a per plate basis. Associations between the presence and absence of accessory genes and phenotype was determined using R package TreeWAS v.1.0 (41).

### RNA-seq analysis

RNA-Seq was performed using data from Lombardi et al. (40). In this analysis, the CLIB214 genome (29) was used instead of CDC317. In brief, reads were aligned the CLIB214 genome using STAR v. 2.7.1a (70) and analysed using deseq2 1.42.1 (71).

### CRISPR-Cas9 gene editing

To test if the DAL4 (CPAR2_103200) Q109* allele observed in some isolate directly affects function, CRISPR-Cas9 gene editing was performed to introduce the stop codon into *C. parapsilosis* FM20 (WT) and to remove the stop codon in *C. parapsilosis* FM33 (DAL4 Q109*). CRISPR-Cas9 gene editing was performed as in Lombardi et al. (72) using the pCP-tRNA plasmid system with guide RNAs designed for the DAL4 gene (Supplementary Table S5). Guide RNAs were designed using EuPaGDT (73). All oligonucleotides (guide RNAs, primers for generating the repair template and primers for screening transformants) are listed in Supplementary Table S5. Plasmids were transformed into *C. parapsilosis* using a lithium acetate based chemical transformation. Successful transformants were screened by incubating on YPD agar supplemented with 200 µg/mL nourseothricin at 30⁰C for 72 h.

### Growth analysis

Isolates were streaked from stock onto YPD agar and were incubated at 30⁰C for 48 h. Single colonies were inoculated into 5 ml YPD broth and incubated at 30⁰C, 200 rpm overnight. Cell pellets were extracted from 1 ml of culture with centrifugation at max speed for 1 min. Cell pellets were washed twice with phosphate-buffered-saline (PBS) solution (BR0014G, Thermo Fisher) and diluted to OD600 values of 0.2 in PBS. The final source plate was created by adding 100 µl of each diluted cell pellet to the wells of a 96-well microtitre plate. Source plates were used to inoculate 4 µl of each isolate into test plates (150 µl ammonium sulphate (0.19% yeast nitrogen base with no amino acids or ammonium sulphate, 0.5% ammonium sulphate, 2% glucose), and 150 µl allantoin (0.19% yeast nitrogen base with no amino acids or ammonium sulphate, 2% glucose, 10mM allantoin)) using a Singer ROTOR HDA robot. Growth was measure using a Synergy H1 microplate reader (Agilent BioTek). Growth was measured using OD600 values every 10 min for 48 h. Area under the curve (AUC) analysis was performed in R v. 4.3.3 using package flux (74).

## Data availability

All sequence data is publicly available at NCBI BioProject PRJNA1173375. Individual accession number for the SRA database are listed in Supplementary Table S1A.

## References

1. Butler G, Rasmussen MD, Lin MF, Santos MA, Sakthikumar S, Munro CA, Rheinbay E, Grabherr M, Forche A, Reedy JL. 2009. Evolution of pathogenicity and sexual reproduction in eight *Candida* genomes. Nature 459:657–662.

2. Santos MA, Tuite MF. 1995. The CUG codon is decoded in vivo as serine and not leucine in *Candida albicans*. Nucl Acids Res 23:1481–1486.

3. Opulente DA, LaBella AL, Harrison M-C, Wolters JF, Liu C, Li Y, Kominek J, Steenwyk JL, Stoneman HR, VanDenAvond J. 2024. Genomic factors shape carbon and nitrogen metabolic niche breadth across Saccharomycotina yeasts. Science 384:eadj4503.

4. Zhai B, Ola M, Rolling T, Tosini NL, Joshowitz S, Littmann ER, Amoretti LA, Fontana E, Wright RJ, Miranda E. 2020. High-resolution mycobiota analysis reveals dynamic intestinal translocation preceding invasive candidiasis. Nat Med 26:59–64.

5. Govrins M, Lass-Flörl C. 2024. *Candida parapsilosis* complex in the clinical setting. Nat Rev Microbiol 22:46–59.

6. Tóth R, Nosek J, Mora-Montes HM, Gabaldon T, Bliss JM, Nosanchuk JD, Turner SA, Butler G, Vágvölgyi C, Gácser A. 2019. *Candida parapsilosis*: from Genes to the Bedside. Clin Microbiol Rev 32.

7. Brassington PJ, Klefisch F-R, Graf B, Pfüller R, Kurzai O, Walther G, Barber AE. 2025. Genomic reconstruction of an azole-resistant *Candida parapsilosis* outbreak and the creation of a multi-locus sequence typing scheme: a retrospective observational and genomic epidemiology study. The Lancet Microbe 6.

8. West PT, Peters SL, Olm MR, Yu FB, Gause H, Lou YC, Firek BA, Baker R, Johnson AD, Morowitz MJ. 2021. Genetic and behavioral adaptation of *Candida parapsilosi*s to the microbiome of hospitalized infants revealed by in situ genomics, transcriptomics, and proteomics. Microbiome 9:1–17.

9. Corzo-Leon DE, Peacock M, Rodriguez-Zulueta P, Salazar-Tamayo GJ, MacCallum DM. 2021. General hospital outbreak of invasive candidiasis due to azole-resistant *Candida parapsilosis* associated with an Erg11 Y132F mutation. Med Mycol 59:664–671.

10. Doorley LA, Barker KS, Zhang Q, Rybak JM, D PR. 2023. Mutations in TAC1 and ERG11 are major drivers of triazole antifungal resistance in clinical isolates of *Candida parapsilosis*. Clin Microbiol Infect 29: 1602.e1–1602.e7.

11. Presente S, Bonnal C, Normand A-C, Gaudonnet Y, Fekkar A, Timsit J-F, Kernéis S. 2023. Hospital clonal outbreak of fluconazole-resistant *Candida parapsilosis* harboring the Y132F ERG11p substitution in a French Intensive Care Unit. Antimicrob Agents and Chemother 67:e01130–22.

12. Branco J, Ryan AP, Pinto ESA, Butler G, Miranda IM, Rodrigues AG. 2022. Clinical azole cross-resistance in *Candida parapsilosis* is related to a novel *MRR1* gain-of-function mutation. Clin Microbiol Infect 28:1655 e5–1655 e8.

13. Berkow EL, Manigaba K, Parker JE, Barker KS, Kelly SL, Rogers PD. 2015. Multidrug Transporters and alterations in sterol biosynthesis contribute to azole antifungal resistance in *Candida parapsilosis*. Antimicrob Agents Chemother 59:5942–50.

14. Bergin SA, Zhao F, Ryan AP, Müller CA, Nieduszynski CA, Zhai B, Rolling T, Hohl TM, Morio F, Scully J. 2022. Systematic analysis of copy number variations in the pathogenic yeast *Candida parapsilosis* identifies a gene amplification in *RTA3* that is associated with drug resistance. MBio 13:e01777–22.

15. Pryszcz LP, Nemeth T, Gacser A, Gabaldon T. 2013. Unexpected genomic variability in clinical and environmental strains of the pathogenic yeast *Candida parapsilosis*. Genome Biol Evol 5:2382–2392.

16. Andreace F, Lechat P, Dufresne Y, Chikhi R. 2023. Comparing methods for constructing and representing human pangenome graphs. Genome Biol 24:274.

17. Contreras-Moreira B, Vinuesa P. 2013. GET_HOMOLOGUES, a versatile software package for scalable and robust microbial pangenome analysis. Appl Environ Microbiol 79:7696–7701.

18. McCarthy CG, Fitzpatrick DA. 2019. Pan-genome analyses of model fungal species. Microb Genom 5:e000243.

19. Jonkheer EM, van Workum D-JM, Sheikhizadeh Anari S, Brankovics B, de Haan JR, Berke L, van der Lee TA, de Ridder D, Smit S. 2022. PanTools v3: functional annotation, classification and phylogenomics. Bioinformatics 38:4403–4405.

20. Barber AE, Sae-Ong T, Kang K, Seelbinder B, Li J, Walther G, Panagiotou G, Kurzai O. 2021. Aspergillus fumigatus pan-genome analysis identifies genetic variants associated with human infection. Nat Microbiol 6:1526–1536.

21. Medini D, Donati C, Tettelin H, Masignani V, Rappuoli R. 2005. The microbial pan-genome. Curr Opin Genet Dev 15:589–594.

22. Hu Z, Wei C, Li Z. 2020. The pangenome: diversity, dynamics and evolution of genomes, in Computational strategies for eukaryotic pangenome analyses. The pangenome: diversity, dynamics and evolution of genomes, pp293–307.

23. Gong Y, Li Y, Liu X, Ma Y, Jiang L. 2023. A review of the pangenome: how it affects our understanding of genomic variation, selection and breeding in domestic animals? J Anim Sci Biotechnol 14:73.

24. Li G, Ji B, Nielsen J. 2019. The pan-genome of *Saccharomyces cerevisiae*. FEMS Yeast Res19:foz064.

25. Peter J, De Chiara M, Friedrich A, Yue J-X, Pflieger D, Bergström A, Sigwalt A, Barre B, Freel K, Llored A. 2018. Genome evolution across 1,011 *Saccharomyces cerevisiae* isolates. Nature 556:339–344.

26. Stamatakis A. 2014. RAxML version 8: a tool for phylogenetic analysis and post-analysis of large phylogenies. Bioinformatics 30:1312–1313.

27. Prjibelski A, Antipov D, Meleshko D, Lapidus A, Korobeynikov A. 2020. Using SPAdes de novo assembler. Curr Protoc Bioinformatics 70:e102.

28. Koren S, Walenz BP, Berlin K, Miller JR, Bergman NH, Phillippy AM. 2017. Canu: scalable and accurate long-read assembly via adaptive k-mer weighting and repeat separation. Genome Res 27:722–736.

29. Cillingová A, Tóth R, Mojáková A, Zeman I, Vrzoňová R, Siváková B, Baráth P, Neboháčová M, Klepcová Z, Brázdovič F. 2022. Transcriptome and proteome profiling reveals complex adaptations of *Candida parapsilosis* cells assimilating hydroxyaromatic carbon sources. PLoS Genetics 18:e1009815.

30. Gabriel L, Brůna T, Hoff KJ, Ebel M, Lomsadze A, Borodovsky M, Stanke M. 2024. BRAKER3: Fully automated genome annotation using RNA-seq and protein evidence with GeneMark-ETP, AUGUSTUS, and TSEBRA. Genome Res 34:769–777.

31. Maguire SL, OhÉigeartaigh SS, Byrne KP, Schröder MS, O’Gaora P, Wolfe KH, Butler G. 2013. Comparative genome analysis and gene finding in Candida species using CGOB. Mol Biol Evol 30:1281–1291.

32. Guida A, Lindstädt C, Maguire SL, Ding C, Higgins DG, Corton NJ, Berriman M, Butler G. 2011. Using RNA-seq to determine the transcriptional landscape and the hypoxic response of the pathogenic yeast *Candida parapsilosis*. BMC Genomics 12:1–14.

33. Blomberg SP, Garland Jr T, Ives AR. 2003. Testing for phylogenetic signal in comparative data: behavioral traits are more labile. Evolution 57:717–745.

34. Münkemüller T, Lavergne S, Bzeznik B, Dray S, Jombart T, Schiffers K, Thuiller W. 2012. How to measure and test phylogenetic signal. Methods Ecol Evol 3:743–756.

35. Revell LJ, Mahler DL, Peres-Neto PR, Redelings BD. 2012. A new phylogenetic method for identifying exceptional phenotypic diversification. Evolution 66:135–146.

36. Yashiroda H, Oguchi T, Yasuda Y, Toh-e A, Kikuchi Y. 1996. Bul1, a new protein that binds to the Rsp5 ubiquitin ligase in *Saccharomyces cerevisiae*. Mol Cell Biol 16:3255–3263.

37. Clark TA, Slavinski SA, Morgan J, Lott T, Arthington-Skaggs BA, Brandt ME, Webb RM, Currier M, Flowers RH, Fridkin SK. 2004. Epidemiologic and molecular characterization of an outbreak of *Candida parapsilosis* bloodstream infections in a community hospital. J Clin Microbiol 42:4468–4472.

38. Kuhn DM, Mukherjee PK, Clark TA, Pujol C, Chandra J, Hajjeh RA, Warnock DW, Soll DR, Ghannoum MA. 2004. *Candida parapsilosis* characterization in an outbreak setting. Emerg Infect Dis 10:1074.

39. Skrzypek MS, Binkley J, Binkley G, Miyasato SR, Simison M, Sherlock G. 2016. The Candida Genome Database (CGD): incorporation of Assembly 22, systematic identifiers and visualization of high throughput sequencing data. Nucl Acids Res 45: D592–D596

40. Lombardi L, Salzberg LI, Cinnéide EÓ, O’Brien C, Morio F, Turner SA, Byrne KP, Butler G. 2024. Alternative sulphur metabolism in the fungal pathogen *Candida parapsilosis*. Nat Commun 15:9190.

41. Collins C, Didelot X. 2018. A phylogenetic method to perform genome-wide association studies in microbes that accounts for population structure and recombination. PLoS Comp Biol 14:e1005958.

42. VanderSluis B, Hess DC, Pesyna C, Krumholz EW, Syed T, Szappanos B, Nislow C, Papp B, Troyanskaya OG, Myers CL. 2014. Broad metabolic sensitivity profiling of a prototrophic yeast deletion collection. Genome Biol 15:1–18.

43. Cooper TG, Gorski M, Turoscy V. 1979. A cluster of three genes responsible for allantoin degradation in *Saccharomyces cerevisiae*. Genetics 92:383–396.

44. Jones P, Binns D, Chang H-Y, Fraser M, Li W, McAnulla C, McWilliam H, Maslen J, Mitchell A, Nuka G. 2014. InterProScan 5: genome-scale protein function classification. Bioinformatics 30:1236–1240.

45. Paluszynski JP, Klassen R, Rohe M, Meinhardt F. 2006. Various cytosine/adenine permease homologues are involved in the toxicity of 5-fluorocytosine in *Saccharomyces cerevisiae*. Yeast 23:707–715.

46. Marcet-Houben M, Alvarado M, Ksiezopolska E, Saus E, de Groot PW, Gabaldón T. 2022. Chromosome-level assemblies from diverse clades reveal limited structural and gene content variation in the genome of *Candida glabrata*. BMC Biol 20:226.

47. Bergin S, Doorley LA, Rybak JM, Wolfe KH, Butler G, Cuomo CA, Rogers PD. 2024. Analysis of clinical Candida parapsilosis isolates reveals copy number variation in key fluconazole resistance genes. Antimicrob Agents Chemother 68:e01619–23.

48. Wisecaver JH, Slot JC, Rokas A. 2014. The evolution of fungal metabolic pathways. PLoS Genet 10:e1004816.

49. Vu D, Groenewald M, Szöke S, Cardinali G, Eberhardt U, Stielow B, De Vries M, Verkleij G, Crous P, Boekhout T. 2016. DNA barcoding analysis of more than 9000 yeast isolates contributes to quantitative thresholds for yeast species and genera delimitation. Stud Mycol 85:91–105.

50. Tavanti A, Davidson AD, Gow NA, Maiden MC, Odds FC. 2005. *Candida orthopsilosis* and *Candida metapsilosis* spp. nov. to replace *Candida parapsilosis* groups II and III. J Clin Microbiol 43:284–292.

51. Pryszcz LP, Nemeth T, Gacser A, Gabaldon T. 2014. Genome comparison of *Candida orthopsilosis* clinical strains reveals the existence of hybrids between two distinct subspecies. Genome Biol Evol 6:1069–1078.

52. Pryszcz LP, Németh T, Saus E, Ksiezopolska E, Hegedűsová E, Nosek J, Wolfe KH, Gacser A, Gabaldón T. 2015. The genomic aftermath of hybridization in the opportunistic pathogen *Candida metapsilosis*. PLoS Genet 11:e1005626.

53. Jiang H, Lei R, Ding S-W, Zhu S. 2014. Skewer: a fast and accurate adapter trimmer for next-generation sequencing paired-end reads. BMC Bioinfo 15:1–12.

54. Hu J, Fan J, Sun Z, Liu S. 2020. NextPolish: a fast and efficient genome polishing tool for long-read assembly. Bioinformatics 36:2253–2255.

55. Cabanettes F, Klopp C. 2018. D-GENIES: dot plot large genomes in an interactive, efficient and simple way. PeerJ 6:e4958.

56. Altschul SF, Madden TL, Schäffer AA, Zhang J, Zhang Z, Miller W, Lipman DJ. 1997. Gapped BLAST and PSI-BLAST: a new generation of protein database search programs. Nucl Acids Res 25:3389–3402.

57. Quinlan AR, Hall IM. 2010. BEDTools: a flexible suite of utilities for comparing genomic features. Bioinformatics 26:841–842.

58. Sayers EW, Cavanaugh M, Clark K, Pruitt KD, Schoch CL, Sherry ST, Karsch-Mizrachi I. 2022. GenBank. Nucl Acids Res 50:D161–D164.

59. Flynn JM, Hubley R, Goubert C, Rosen J, Clark AG, Feschotte C, Smit AF. 2020. RepeatModeler2 for automated genomic discovery of transposable element families. Proc Natl Acad Sci 117:9451–9457.

60. Katoh K, Misawa K, Kuma Ki, Miyata T. 2002. MAFFT: a novel method for rapid multiple sequence alignment based on fast Fourier transform. Nucl Acids Res 30:3059–3066.

61. Eddy S. 1992. HMMER user’s guide. Department of Genetics, Washington University School of Medicine 2:13.

62. Steenwyk JL, Rokas A. 2021. Orthofisher: a broadly applicable tool for automated gene identification and retrieval. G3 11:jkab250.

63. Rausch T, Zichner T, Schlattl A, Stütz AM, Benes V, Korbel JO. 2012. DELLY: structural variant discovery by integrated paired-end and split-read analysis. Bioinformatics 28:i333–i339.

64. Li H. 2013. Aligning sequence reads, clone sequences and assembly contigs with BWA-MEM. arXiv preprint arXiv:13033997.

65. Danecek P, Bonfield JK, Liddle J, Marshall J, Ohan V, Pollard MO, Whitwham A, Keane T, McCarthy SA, Davies RM. 2021. Twelve years of SAMtools and BCFtools. Gigascience 10:giab008.

66. Van der Auwera GA, Carneiro MO, Hartl C, Poplin R, Del Angel G, Levy-Moonshine A, Jordan T, Shakir K, Roazen D, Thibault J. 2013. From FastQ data to high-confidence variant calls: the genome analysis toolkit best practices pipeline. Curr Protoc Bioinformatics 43:11.10. 1–11.10. 33.

67. Richter F, Morton SU, Qi H, Kitaygorodsky A, Wang J, Homsy J, DePalma S, Patel N, Gelb BD, Seidman JG. 2020. Whole genome de novo variant identification with FreeBayes and neural network approaches. BioRxiv:2020.03.24.994160.

68. Vaser R, Adusumalli S, Leng SN, Sikic M, Ng PC. 2016. SIFT missense predictions for genomes. Nat Protoc 11:1–9.

69. Virtanen P, Gommers R, Oliphant TE, Haberland M, Reddy T, Cournapeau D, Burovski E, Peterson P, Weckesser W, Bright J. 2020. SciPy 1.0: fundamental algorithms for scientific computing in Python. Nat Method 17:261–272.

70. Dobin A, Davis CA, Schlesinger F, Drenkow J, Zaleski C, Jha S, Batut P, Chaisson M, Gingeras TR. 2013. STAR: ultrafast universal RNA-seq aligner. Bioinformatics 29:15–21.

71. Love M, Anders S, Huber W. 2014. Differential analysis of count data–the DESeq2 package. Genome Biol 15:10–1186.

72. Lombardi L, Oliveira-Pacheco J, Butler G. 2019. Plasmid-based CRISPR-Cas9 gene editing in multiple Candida species. MSphere 4:10.1128/msphere.00125-19.

73. Peng D, Tarleton R. 2015. EuPaGDT: a web tool tailored to design CRISPR guide RNAs for eukaryotic pathogens. Microb Genom 1:e000033.

74. Jurasinski G, Koebsch F, Guenther A, Beetz S, Jurasinski MG. 2014. Package ‘flux’. Flux rate calculation from dynamic closed chamber measurement: R.

